# Orbitofrontal-striatal potentiation underlies stimulant induced hyperactivity

**DOI:** 10.1101/802900

**Authors:** Sebastiano Bariselli, Nanami Miyazaki, Alexxai Kravitz

**Author notes:** **Corresponding author:**Alexxai V. Kravitz 425 S Euclid Ave, CSRB 5536 Saint Louis, MO 63110.

## Abstract

Stimulants are one of the most widely prescribed classes of pharmaceuticals, but it is unclear which brain pathways underlie their therapeutic and adverse actions. Here, with real-time monitoring of circuit plasticity, we demonstrate that psychostimulants strengthen orbitofrontal (OFC) to dorsomedial striatum (DMS) pathway synapse*s*, and increase striatal output in awake mice. *In vivo* high-frequency stimulation of OFC-DMS pathway blocked stimulant-induced potentiation and the expression of locomotor sensitization, thereby directly linking OFC-DMS plasticity to hyperactivity.

## Main text

Psychomotor stimulants are widely prescribed drugs for neurodevelopmental diseases, including attention-deficit hyperactivity disorder (ADHD)^1^. However, their use is also associated with adverse effects, such as hyperactivity, cognitive impairments, and sensitization and tolerance, which eventually result in substance abuse^1^. Uncovering the brain pathways underlying their adverse actions is fundamental to develop refined therapeutic strategies. *In vitro* work has been instrumental for linking plasticity in the orbitofrontal cortex (OFC) to dorsomedial striatum (DMS) projection to compulsive behavior^2,3^. However, the slice preparation does not allow for the examination of these adaptations as they occur, nor directly link such changes to ongoing behavior. To overcome these limitations, we utilized an awake *in vivo* preparation to demonstrate that OFC-DMS potentiation underlies stimulant-induced hyperactivity. Moreover, we found that high-frequency stimulation (HFS) blocks the stimulant-induced OFC-DMS potentiation and the expression of locomotor sensitization. Thus, our data suggest that modulating the function of the orbitofrontal-striatal pathway might be a potential strategy to counteract, at least some of, the adverse effects of psychostimulants.

To determine which striatal subdivisions were most activated by psychostimulants, we analysed the expression of phospho-c-Fos (Ser32) following an intraperitoneal (i.p.) injection of cocaine (20 mg/kg, Figure 1a). Similar to previous evidence^4^, cocaine increased phospho-c-Fos immunoreactivity throughout dorsal and ventral striatum (Figure 1b) with the strongest effect in the dorsomedial striatum (Figure 1b,c). Cocaine increases striatal dopamine *in vivo*^5^ that has been shown, both *ex vivo*^6^ and in anaesthetized animals^7^, to modulate cortico-striatal plasticity. Thus, we hypothesized that this activation may be associated with a potentiation of OFC-DMS efficacy, either by enhancing synaptic connections^2^ or promoting “up-states” in striatal neurons^8^. To test whether OFC-DMS inputs were potentiated during psychomotor stimulant exposure, we expressed the blue light activated opsin Chronos^9^ in the OFC (Figure 1d) and measured cortical stimulation-evoked changes in striatal local field potentials (LFPs), a measure of cortical-input efficacy^10,11^. Mice received an i.p. injection of either cocaine, d-amphetamine (d-amph), or saline for controls (cSAL and dSAL, Figure 1e). We probed the OFC continually with brief (15 msec) pulses at 0.2 Hz and found that cocaine and d-amph (3 mg/Kg i.p.) both increased locomotor activity (Supplementary Figure 1a,b) and the amplitude of OFC-evoked (OFCe) striatal LFP responses compared to their respective saline controls (Figure 1f-h, Supplementary figure 1c-e). We conclude that both cocaine and amphetamine increase striatal responsivity to OFC input.

**Figure 1.**
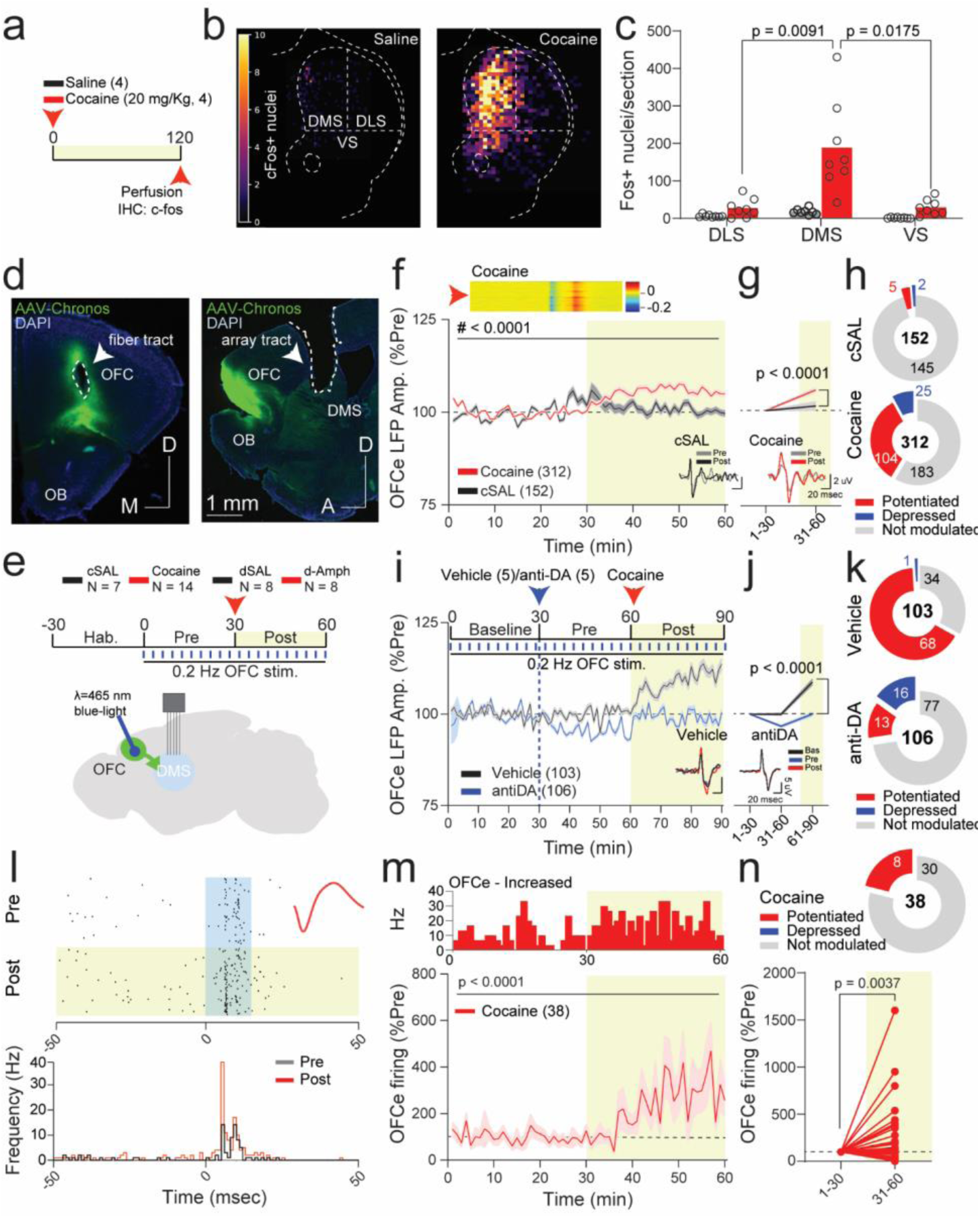
Hyperdopaminergic states enhance OFC-DMS synaptic efficacy. (a) Experimental design of c-fos experiments. (b) Brain slice heat-maps of c-fos positive nuclei within striatal sub-regions (DLS: dorsolateral, DMS: dorsomedial, VS: ventral). (c) Scatter-plot of averaged c-fos positive nuclei across striatal subregions (RM two-way ANOVA; subregion main effect: F_(1.127,15.78)_ = 19.27, p = 0.0003; drug main effect: F_(1,14)_ = 17.13, p = 0.0010; subregion Χ drug interaction: F_(2,28)_ = 13.63, p < 0.0001; followed by Bonferroni post-hoc test, DLS vs DMS: t_(7)_ = 4.433; DMS vs VS: t_(7)_ = 3.909). (d) Representative images of Chronos expression in OFC (OB: olfactory bulb). (e) Experimental and brain schematics for electrophysiology experiments. (f) Time-course of normalized OFCe LFP amplitude (two-way ANOVA; time main effect: F_(59,27720)_ = 13.18, p < 0.0001; drug main effect: F_(1,27720)_ = 225.7, p < 0.0001; time Χ drug interaction: F_(59, 27720)_ = 10.27, p < 0.0001). (g) Binned OFCe LFP amplitude (two-way ANOVA; time main effect: F_(1,924)_ = 35.11, p < 0.0001; drug main effect: F_(1,924)_ = 14.27, p = 0.0002, time Χ drug interaction: F_(1,924)_ = 14.27, p = 0.0002; followed by Bonferroni post-hoc test; Pre: cSAL vs cocaine: t_(924)_ = 0, p > 0.9999; post: t_(924)_ = 5.342). (h) Pie-charts reporting the number of significantly potentiated, depressed or non-significantly modulated OFCe LFP responses. (i) Top: experimental schematics of D1R and D2R antagonist pre-treatment. Bottom: time-course of normalized OFCe LFP amplitude (RM two-way ANOVA; time main effect F_(3.568,738.7)_ = 19.79, p < 0.0001, drug main effect: F_(1,107)_ = 133.8, p < 0.0001, time Χ drug interaction: F_(89,18423)_ = 14.18, p < 0.0001) (j) Binned OFCe LFP responses (RM two-way ANOVA, time main effect: F_(1.277,264.3)_ = 150.4, p < 0.0001; drug main effect: F_(1,207)_ = 122.3, p < 0.0001; time Χ drug: F_(2,414)_ = 75.39, p < 0.0001; followed by Bonferroni post-hoc test antiDA vs Veh; 1-30: t_(621)_ = 0, p > 0.9999; 31-60: t_(621)_ = 5.923; 61-90: t_(621)_ = 15.87). (k) Pie-charts reporting the number of non-modulated, potentiated or depressed OFCe LFP responses. (l) Representative raster-plot and peri-stimulation histogram of OFCe responses. (m) Time-course of normalized OFC-stimulation evoked firing rate (RM one-way ANOVA, F_(59,2183)_ = 3.056). (n) Top: pie-charts reporting the number of OFCe firing responses. Bottom: binned OFCe firing frequency in response to cocaine (Wilcoxon matched pairs signed rank test: W = 392.0). Data are represented as single points and/or as average ± SEM. N indicates number of mice or OFCe LFPs and units included in the analysis.

Considering that cocaine blocks dopamine (DAT) but also other monoaminergic transporters^12^, we sought to isolate the contribution of dopamine to the potentiation of OFC-DMS connection strength. We first assessed whether an i.p. injection of a more selective DAT blocker^13^, GBR13069, recapitulated the effects induced by psychostimulant exposure. GBR13069 increased both locomotor activity (Supplementary figure 1f) and OFCe LFP responses (Supplementary figure 1g-i) similar to cocaine and d-amphetamine (Supplementary figure 1j). We also tested whether blocking dopamine receptors with a mixture of antagonists for D1R (SCH23390) and D2R (sulpiride) would prevent the cocaine-induced increase in OFCe LFP amplitude (Figure 1i). Injecting this dopamine antagonist mix (anti-DA) caused a slight depression in OFCe LFP amplitude (Figure 1i,j), demonstrating that physiological dopamine receptor activation controls orbitofrontal-striatal pathway input strength. Pre-treatment with anti-DA also blunted both the cocaine-induced hyperlocomotion (Supplementary figure 1k) and the increase of OFCe LFP amplitude (Figure 1i-k), even when normalizing for the anti-DA mediated reduction in OFCe LFPs (Supplementary figure 1l). Thus, we conclude that psychomotor stimulants increase OFC-DMS input strength in a dopamine-dependent manner. Next, to investigate whether changes in input transmission were associated with alterations in striatal neuronal firing, we examined the effects of cocaine on OFCe firing of striatal multi- and single units. Forty-four of 197 (22%) recorded striatal units were modulated by OFC stimulation, and most (38 of 44, 86%) responsive units were excited, consistent with a glutamatergic input from OFC (Supplementary figure 1m-o). Similar to its effect on OFCe LFPs, cocaine also potentiated OFCe spiking in these units (Figure 1l-n). We next analysed DMS firing rates after exposure to psychostimulants (Figure 2a, Supplementary figure 2a). In agreement with our c-fos expression data and OFCe potentiation of LFPs and spiking, both cocaine and amphetamine increased overall striatal multi-unit firing rates (Figure 2b,c and Supplementary figure 2b). This increase was also observed when we limited our analysis of firing rates to well-isolated single MSNs (time peak-to-valley > 350 μsec; Supplementary figure 2a,c-i). We conclude that psychomotor stimulants potentiate OFC-DMS inputs, increase synaptic efficacy, and enhance spiking output of the DMS in awake mice.

**Figure 2.**
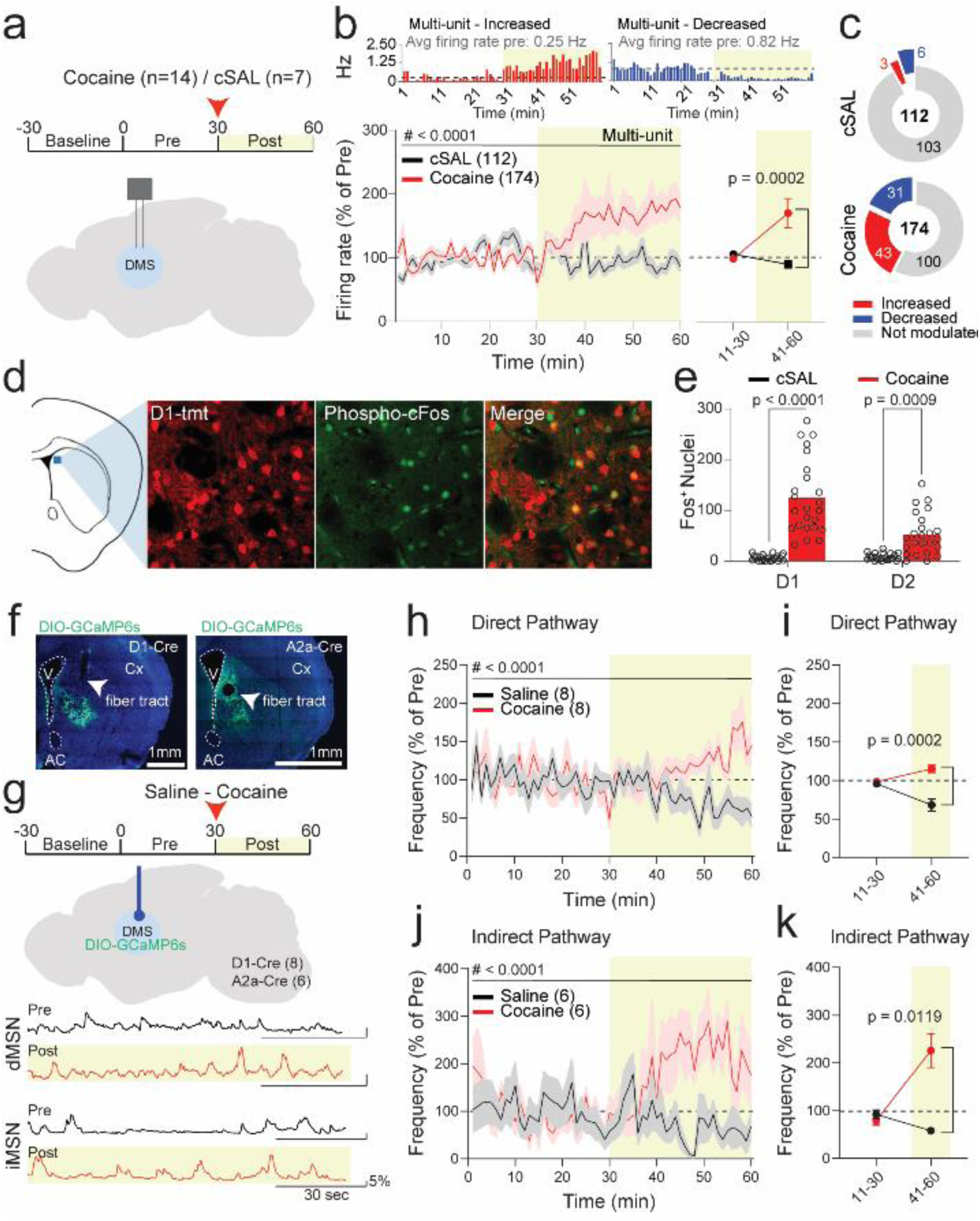
Cocaine increases both direct and indirect pathway neuronal activity. (a) Experimental and brain schematic. (b) Time-course of normalized multi-unit activity (RM two-way ANOVA; time main effect: F_(3.063,869.8)_ = 3.118, p = 0.0246; drug main effect: F_(1,284)_ = 6.076, p = 0.0143, time Χ drug interaction: F_(59,16756)_ = 4.180, p < 0.0001) and binned analysis of normalized multi-unit activity (RM two-way ANOVA; time main effect: F_(1,284)_ = 3.732, p = 0.0544; drug main effect: F_(1,284)_ = 6.511, p = 0.0112; time Χ drug interaction: F_(1,284)_ = 9.185, p = 0.0027, followed by Bonferroni post-hoc test cSAL vs COC; 11-30: t_(568)_ = 0.3359, p > 0.9999; 41-60: t_(568.0)_ = 3.947). Top: example rate histograms. (c) Pie-chart with modulated or not modulated multi-units. (d) Representative images of phospho-c-fos staining: D1-tmt (D1-tomato; red) and phospho-c-fos (green). (e) Phospho-c-fos quantification in direct or indirect pathway neurons of DMS (two-way ANOVA; cell type main effect: F_(1,92)_ = 16.72, p < 0.0001; drug main effect: F_(1,92)_ = 88.83, p < 0.0001; cell type Χ drug interaction: F_(1,92)_ = 18.44, p < 0.0001; followed by Bonferroni post-hoc test cSAL vs COC; D1: t_(92)_ = 9.701; D2: t_(92)_ = 3.628). (f) Representative images of GCaMP6s expression in DMS (V: ventricle, AC: anterior commissure, Cx: cortex). (g) Experimental and brain schematics of fiber photometry experiments with example photometry traces. (h) Norm. frequency of direct pathway calcium events (RM two-way ANOVA by both factors; time main effect: F_(59,413)_ = 1.063, p = 0.3585; drug main effect: F_(1,7)_ = 20.37, p = 0.0028; time Χ drug interaction: F_(59,413)_ = 2.891, p < 0.0001). (i) Binned norm. frequency of direct pathway calcium events (RM two-way ANOVA by both factors; time main effect: F_(1,7)_ = 0.9040, p = 0.3734; drug main effect: F_(1,7)_ = 53.77, p = 0.0002, time Χ drug interaction: F_(1,7)_ = 72.32, p < 0.0001, followed by Bonferroni post-hoc test Saline vs Cocaine; 11-30: t_(7)_ = 1.539, p = 0.3354; 31-60: t_(7)_ = 8.137). (j) Norm. frequency of indirect pathway calcium events (RM two-way ANOVA by both factors; time main effect: F_(59,295)_ = 1.118, p = 0.2729; drug main effect: F_(1,5)_ = 21.45, p = 0.0057; time Χ drug interaction: F_(59,295)_ = 2.5, p < 0.0001). (k) Binned frequency of indirect pathway calcium events (RM two-way ANOVA by both factors; time main effect: F_(1,5)_ = 9.215, p = 0.0289; drug main effect: F_(1,5)_ = 13.48, p = 0.0144; time Χ drug interaction: F_(1,5)_ = 25.37, p = 0.0040; followed by Bonferroni post-hoc test Saline vs Cocaine; 11-30: t_(5)_ = 0.9309, p = 0.7893; 31-60: t_(5)_ = 4.583). Data are represented as single points and/or as mean ± SEM. N indicates number of multi-units or animals included in the analysis.

Previous studies found bidirectional, or inhibitory, changes of striatal activity during hyperdopaminergic states^14–16^, depending on the experimental paradigms and/or striatal region. Dopamine modulates striatal output pathway activity *via* post-synaptic dopamine 1 (D1) and D2 receptors which increase and decrease medium spiny neuron (MSN) excitability, respectively^17^. However, it remains unclear whether cocaine increases and decreases the activity of D1R-expressing MSNs (direct pathway, dMSNs) and D2R-expressing MSNs (indirect pathway, iMSNs)^14–16^, respectively. To test this, we first analysed expression of phosho-c-Fos in dMSNs or iMSNs, differentiated by genetic expression of a fluorescent label. Similarly to previous reports, phospho-c-Fos expression increased in both populations (Figure 2d,e), suggesting that cocaine activates both pathways^4^. To assess this question in real time, D1Cre and A2aCre mice were infected with a Cre-dependent DIO-GCaMP6s virus (Figure 2f) and implanted with an optic fiber in the DMS to record calcium bulk signal from each pathway in this region (Figure 2g). D1Cre and A2aCre mice had similar frequencies of calcium events during pre-cocaine period (Supplementary figure 2j) and comparable hyper-locomotor responses to cocaine (Supplementary figure 2k). Following an i.p, injection of cocaine, population calcium transient frequency increased in both direct (Figure 2h,i) and indirect pathways (Figure 2j,k). Our data indicate that the cocaine-induced potentiation of OFC-striatal synaptic efficacy is accompanied by activation of both striatal output pathways.

As cocaine potentiated OFC-DMS inputs, we asked whether plasticity-inducing stimulation protocols could be identified that would depress this input and block cocaine-induced behaviour, building on similar approaches used in prior research^18–21^. We tested the effect of multiple stimulation protocols based on those that induce plasticity *ex vivo* (Figure 3a, Supplementary figure 3a,b). Testing these plasticity protocols in awake mice allowed us to monitor circuit plasticity as it occurs, which is critical for understanding how they affect these circuits *in vivo*. Thus, we first investigated the consequences of low-frequency (LFS; 5 Hz for 15 min) and theta-burst (TBS; 10 repetitions of 40 stimulations organized in trains of 50Hz every 10.5Hz) stimulation of OFC on the magnitude of OFCe LFP responses in DMS, since similar protocols cause long-term depression^22^ or potentiation^23^, respectively, of cortico-striatal synapses in *ex vivo* preparations. Surprisingly, both LFS and TBS failed to induce plasticity in our awake preparation (Supplementary figure 3c-h). We next delivered a high-frequency stimulation (sHFS: short high frequency stimulation; 60Hz for 5 min), which also depresses excitatory synapses in striatal slices^24^. This protocol decreased OFCe LFP amplitude in the awake preparation (Supplementary figure 3i-k).

**Figure 3.**
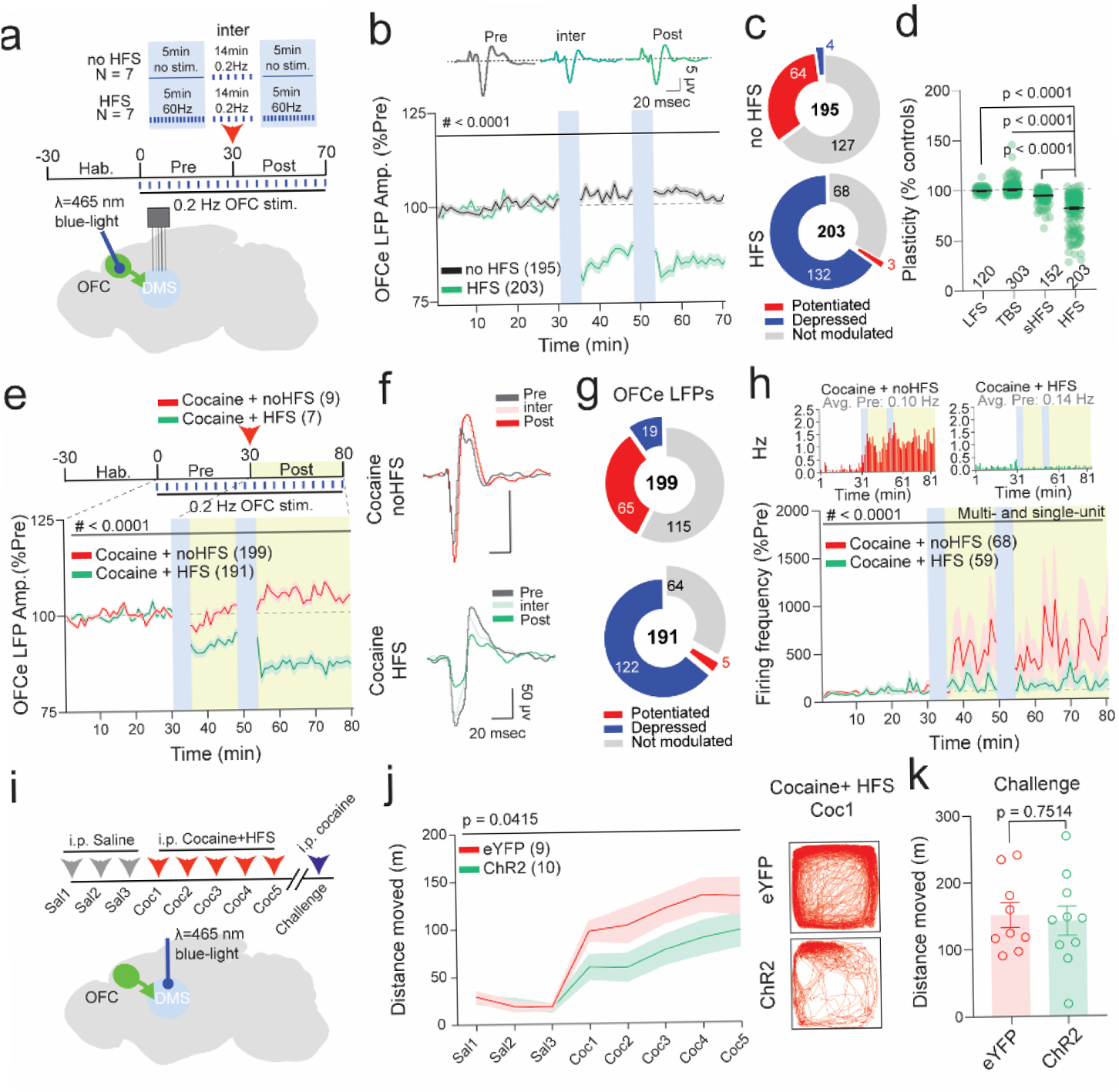
HFS weakens OFC-DMS pathway and attenuates cocaine-induced hyperlocomotion. (a) HFS protocol and brain schematics. (b) Time-course of norm. OFCe LFP (RM two-way ANOVA; time main effect: F_(73,28908)_ = 39.19, p < 0.0001; protocol main effect: F_(1,396)_ = 205.2, p < 0.0001; time Χ protocol interaction: F_(73,28908)_ = 89.20, p < 0.0001). (c) Pie-charts reporting the number of non-modulated, potentiated or depressed OFCe LFP responses. (d) Scatter-plot reporting the change in OFCe LFP responses relative to controls for different stimulation protocols (one-way ANOVA; F_(3,774)_ = 181.7, p < 0.0001 followed by Dunnett’s test; HFS vs LFS: q_(774)_ = 16.72; HFS vs TBS: q_(774)_ = 22.25; HFS vs sHFS: q_(774)_ = 12.42). (e) Time-course of norm. OFCe LFP responses (RM two-way ANOVA; time main effect: F_(69,26772)_ = 23.08, p < 0.0001; treatment main effect: F_(1,388)_ = 130.3, p < 0.0001; time Χ treatment interaction: F_(69,26772)_ = 71.29, p < 0.0001). (f) Representative OFCe LFP traces. (g) Pie-charts reporting the number of non-modulated, potentiated or depressed OFCe LFP responses. (h) Time-course of norm. firing frequency of multi- and single-units (RM two-way ANOVA; time main effect: F_(73,9125)_ = 2.499, p < 0.0001; treatment main effect: F_(1,125)_ = 2.106, p = 0.1492; time Χ treatment interaction: F_(73,9125)_ = 1.756, p < 0.0001). Top: representative rate histograms. (i) Experimental schematics. (j) Distance moved during cocaine sensitization paradigm (RM two-way ANOVA; time main effect: F_(7,119)_ = 28.91, p < 0.0001; virus main effect: F_(1,17)_ = 4.861, p = 0.0415; time Χ virus interaction: F_(7,119)_ = 2.009, p = 0.0595). (k) Distance moved upon cocaine challenge (unpaired t-test: t_(17)_ = 0.3219). Data are represented as single points and/or as mean ± SEM. N indicates number of LFPs, multi-units or animals included in the analysis.

Doubling the sHFS protocol (HFS: 2 sHFS stimulation interleaved by a pause of 14 minutes) caused an even larger depression of OFCe LFP responses (Figure 3b-d, Supplementary figure 3l). We therefore tested whether this HFS protocol could block the cocaine-induced potentiation of the OFC-DMS pathway. In control recordings (without HFS), cocaine again increased OFCe LFP amplitude, but when HFS was applied to the OFC immediately after the cocaine injection it induced a marked depression of OFC-DMS pathway that over-powered the cocaine-induced potentiation (Figure 3e-g). Importantly, HFS also blocked the cocaine-mediated increase in the average firing rate of striatal units (Figure 3h), suggesting a causal link between the cocaine-mediated potentiation of the OFC-DMS pathway and increases in neuronal activity in the striatum.

Finally, we examined the consequences of attenuating cortico-striatal drive on the sensitization of psychomotor responses to cocaine. For 5 consecutive days, mice bilaterally expressing either ChR2 or eYFP in the OFC and implanted with optic fibers in DMS (Figure 3i) received i.p. injections of cocaine (20 mg/Kg) immediately followed by HFS. At the first cocaine exposure (Coc1), HFS blocked hyperactivity in ChR2-, but not eYFP-expressing mice (Supplementary Figure 3m). Attenuation of locomotor responses was also observed over the next 4 consecutive cocaine injections (Figure 3j). To test whether HFS at OFC-DMS was masking the expression of cocaine sensitization, these same animals received a cocaine challenge injection (20 mg/kg, no HFS) after 10 days of withdrawal from cocaine. Without HFS, ChR2- and eYFP-expressing mice displayed similar locomotor responses to this cocaine challenge (Figure 3k). Thus, we conclude that HFS at OFC-DMS pathway did not prevent the acquisition of sensitization but interfered with the expression of psychomotor responses to cocaine.

Here, we adopted an optogenetic-based method to assess input efficacy at cortico-striatal projections *in vivo*, in awake mice, and used it to uncover a role for OFC-DMS pathway in mediating some of the adverse actions of psychomotor stimulants. By monitoring the strength of cortico-striatal inputs in real time, we found that hyperdopaminergic states acutely enhance OFC-DMS input efficacy, which may also relate to similar increases in functional connectivity in chronic cocaine users^25,26^. We further found that HFS of the OFC counteracts the cocaine-induced increase in OFC-DMS input efficacy, and blocks cocaine-induced increases in striatal spiking and locomotion. While cocaine induces plasticity at multiple locations in the brain^27^, stimulation of OFC terminals in the DMS was sufficient to attenuate hyper-locomotion and interfere with the expression of behavioural sensitization to cocaine. Monitoring the strength of the OFC-DMS connection in real time, as we did here, can link plasticity in specific brain circuits in animals to changes in connectivity observed in clinical studies^25,26^. Thus, this approach may be used in multiple circuits to prevent or rescue aberrant forms of plasticity, induced by drugs or diseases, *via* deep-brain stimulation (DBS) or trans-cranial magnetic stimulation (TMS) protocols in both rodents and humans^18–20^.

---

**Supplementary figure 1.**
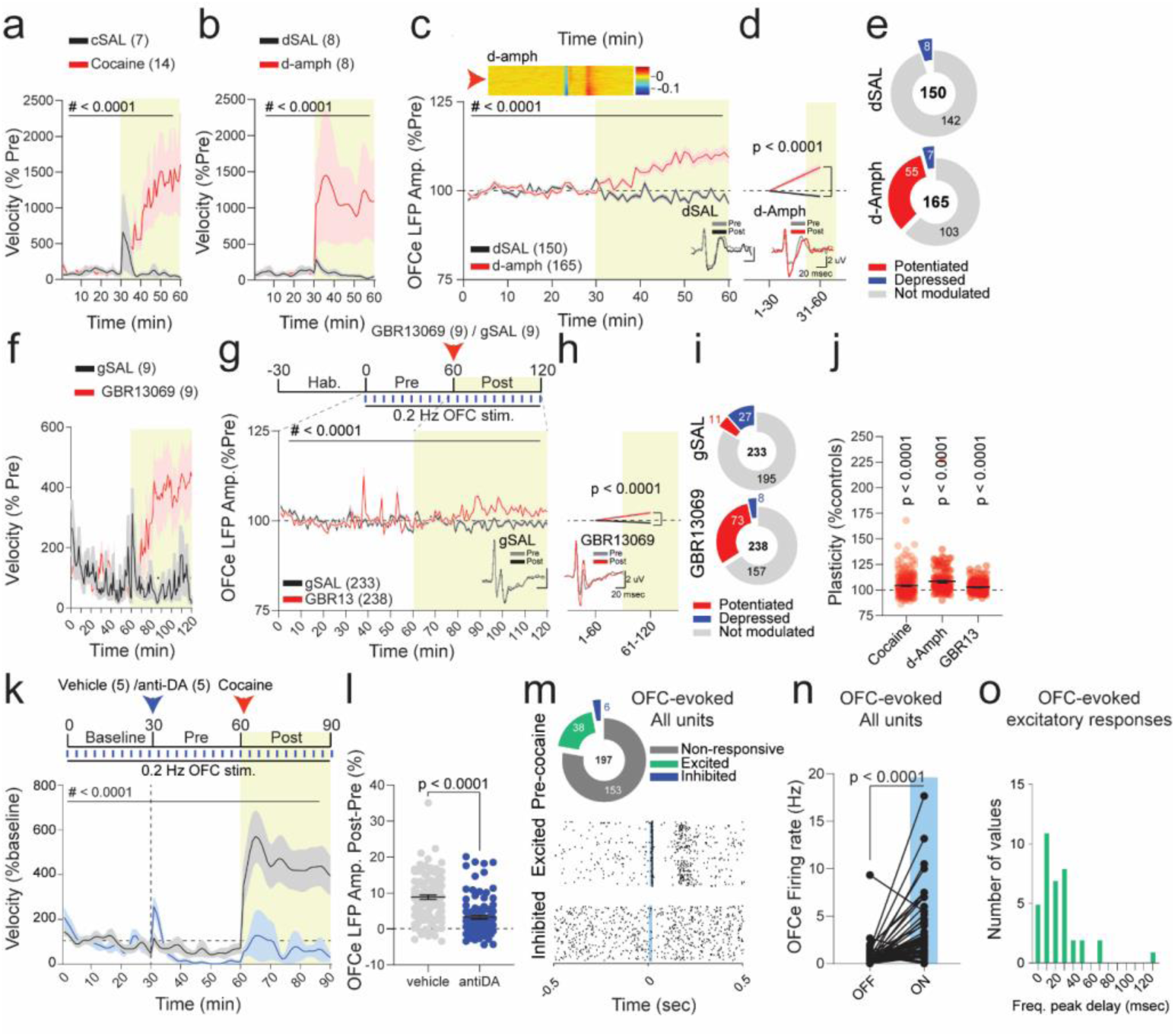
Effects of psychostimulants on velocity, OFCe LFPs and OFCe firing. (a) Time-course of norm. velocity for cocaine and saline (RM two-way ANOVA; time main effect: F_(59,1121)_ = 2.765, p < 0.0001; drug main effect: F_(1,19)_ = 3.829, p = 0.0652; time Χ drug interaction: F_(59,1121)_ = 3.044, p < 0.0001). (b) Time-course of norm. velocity for d-amph or saline (RM two-way ANOVA, time main effect: F_(1.064,14.90)_ = 2.685, p = 0.1209; drug main effect: F_(1,14)_ = 3.039, p = 0.1032; time Χ drug interaction: F_(59,826)_ = 2.753, p < 0.0001). (c) Time-course of normalized OFCe LFP responses following saline or d-amph i.p. injections (two-way ANOVA; time main effect: F_(59,18780)_ = 10.90, p < 0.0001; drug main effect: F_(1,18780)_ = 744.80, p < 0.0001; time Χ drug interaction: F_(59,18780)_ = 18.63, p < 0.0001). (d) Binned OFCe LFP amplitude upon d-SAL or d-amph injection (two-way ANOVA; time main effect: F_(1,626)_ = 21.20, p < 0.0001; drug main effect: F_(1,626)_ = 54.76, p < 0.0001; time Χ drug interaction: F_(1,626)_ = 54.76, p < 0.0001, followed by Bonferroni post-hoc test dSAL vs d-Amph; 1-30: t_(626)_ = 0, p > 0.9999; 31-60: t_(626)_ = 10.47). (e) Pie-charts with numbers of non-significantly modulated, potentiated or depressed OFCe LFP responses. (f) Time-course of norm. velocity upon GBR13069 or saline i.p. injection (RM two-way ANOVA; time main effect: F_(4.840,77.44)_ = 3.588, p = 0.0062; drug main effect: F_(1,16)_ = 9.044, p = 0.0084; time Χ drug interaction: F_(119,1904)_ = 4.656, p < 0.0001). (g) Time-course of normalized OFCe LFP responses upon saline or GBR13069 i.p. injection (RM two-way ANOVA; time main effect: F_(119,56280)_ = 13.83, p < 0.0001; drug main effect: F_(1,56280)_ = 415.1, p < 0.0001; time Χ drug interaction: F_(119,56280)_ = 16.61, p < 0.0001). (h) Binned OFCe LFP amplitude upon saline or GBR13069 i.p. injection (two-way ANOVA; time main effect: F_(1,938)_ = 12.42, p = 0.0004; drug main effect: F_(1,938)_ = 50.17, p < 0.0001; time Χ drug interaction: F_(1,938)_ = 50.17, p < 0.0001; followed by Bonferroni post-hoc test gSAL vs GBR13; 1-30: t_(938)_ = 0, p> 0.9999; 31-60: t_(938)_ = 10.02). (i) Pie-charts reporting the number of non-modulated, potentiated or depressed OFCe LFP responses. (j) Raster plot of % change in OFCe LFP responses upon application of cocaine, d-Amph and GBR13069 relative to their respective controls (one-way ANOVA, F_(2,712)_ = 12.76, p < 0.0001, followed by one-sided t-test to 100%, cocaine: t_(311)_ = 9.469; d-amph: t_(164)_ = 6.421, GBR13: t_(237)_ = 6.120). (k) Time-course of norm. velocity in mice exposed to cocaine and pre-treated with either vehicle or anti-DA (RM two-way ANOVA by both factors; time main effect: F_(89,356)_ = 11.27, p < 0.0001; treatment main effect: F_(1,4)_ = 14.47, p = 0.0190; time Χ treatment interaction: F_(89,356)_ = 8.964, p < 0.0001). (l) Raster-plot of OFCe LFP amplitude difference between Pre and Post period, expressed as percentage of baseline (Mann-Whitney U = 2627). (m) Pie-charts with example raster-plots of striatal units excited or inhibited by OFC stimulation. (n) OFCe firing rate before (OFF) and after OFC stimulation (ON; Wilcoxon matched-pairs signed rank test: W = 823.0). (o) Frequency distribution of peak latency of OFC evoked responses. Data are represented as single points and/or as mean ± SEM. N indicates number of animals, multi-units or OFCe LFPs included in the analysis.

**Supplementary figure 2.**
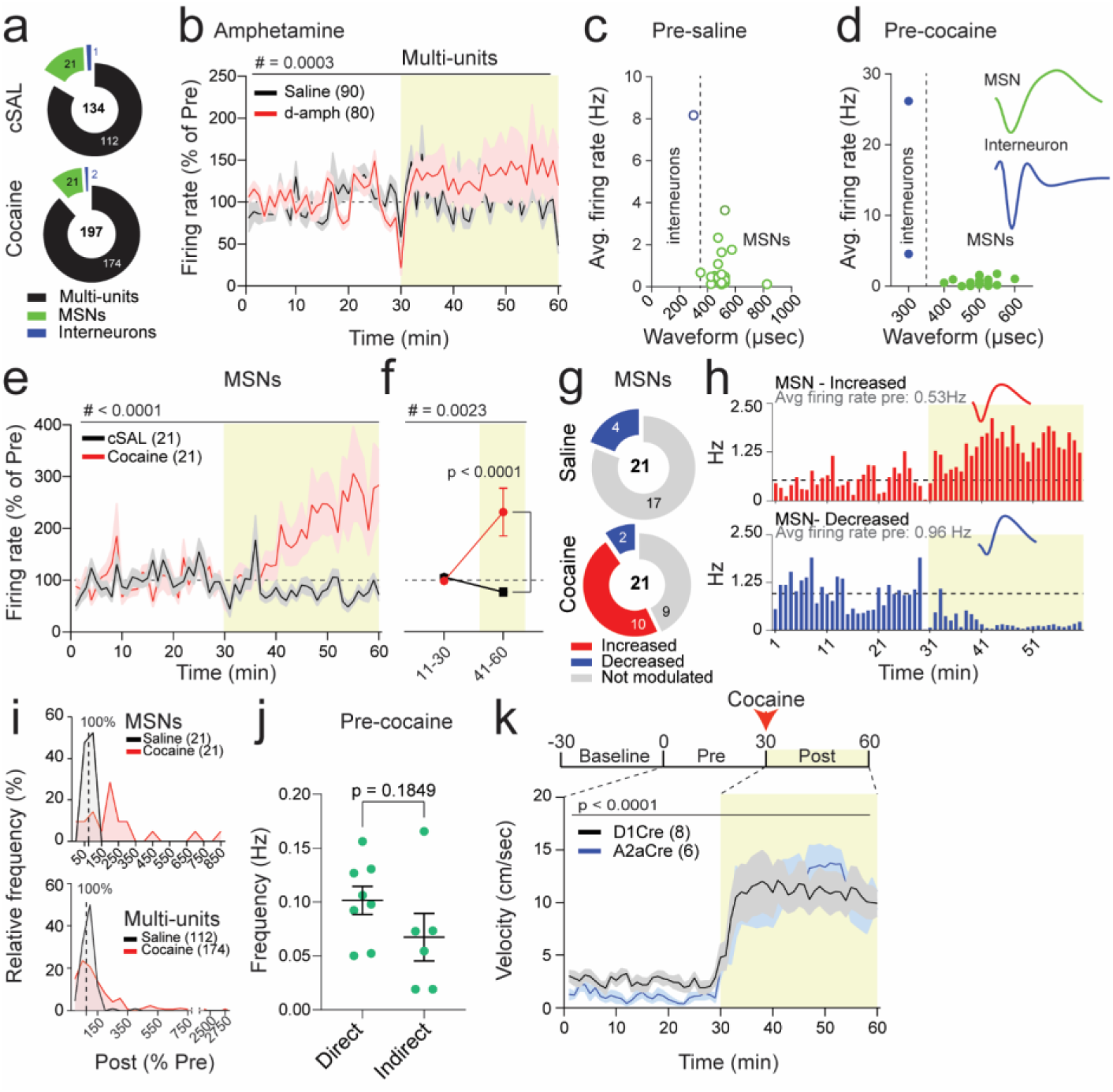
Characterization of multi-unit and MSN activity in DMS. (a) Pie-charts reporting the number of units recorded from saline or cocaine i.p. injected animals. (b) Time-course of norm. multi-unit activity upon saline or d-amph injection (RM two-way ANOVA; time main effect: F_(5.021,843.5)_ = 2.394, p = 0.0359; drug main effect: F_(1,168)_ = 1.091, p = 0.2977; time Χ drug interaction: F_(59,9912)_ = 1.762, p = 0.0003). (c) Raster-plot of average firing and waveform duration of striatal units recorded before saline. (d) Raster-plot of average firing and waveform duration of striatal units recorded before cocaine. (e) Normalized firing rate of MSN activity following saline or cocaine i.p. injection (RM two-way ANOVA; time main effect: F_(4.656,186.2)_ = 2.889, p = 0.0179; drug main effect: F_(1,40)_ = 11.07, p = 0.0019; time Χ drug interaction: F_(59,2360)_ = 4.134, p < 0.0001). (f) Binned norm. firing rate of MSN upon saline or cocaine i.p. injection (RM two-way ANOVA; time main effect: F_(1,40)_ = 4.397, p = 0.0424; drug main effect: F_(1,40)_ = 11.45, p = 0.0016; time Χ drug interaction: F_(1,40)_ = 10.64, p = 0.0023; followed by Bonferroni post-hoc test cSAL vs Cocaine; 11-30: t_(80)_ = 0.2213, p > 0.9999; 41-60: t_(80)_ = 4.684). (g) Pie-charts reporting the number of MSNs with non-modulated, increased or decreased activity upon saline or cocaine i.p. injection. (h) Example rate histograms of MSN increasing or decreasing their activity upon cocaine i.p. injection. (i) Relative frequency distribution of increased or decreased MSN and multi-unit activity upon saline or cocaine injection. (j) Raster plot of calcium population event frequency from direct and indirect pathway before cocaine injection (unpaired t-test, t_(12)_ = 1.407). (k) Time-course of velocity for D1-Cre and A2a-Cre mice injected with cocaine (RM two-way ANOVA; time main effect: F_(2.809,33.71)_ = 31.11, p < 0.0001; genotype main effect: F_(1,12)_ = 0.2283, p = 0.6414; time Χ genotype interaction: F_(59,708)_ = 0.8194, p = 0.8304). Data are represented as single points and/or as mean ± SEM. N indicates number of units or animals included in the analysis.

**Supplementary figure 3.**
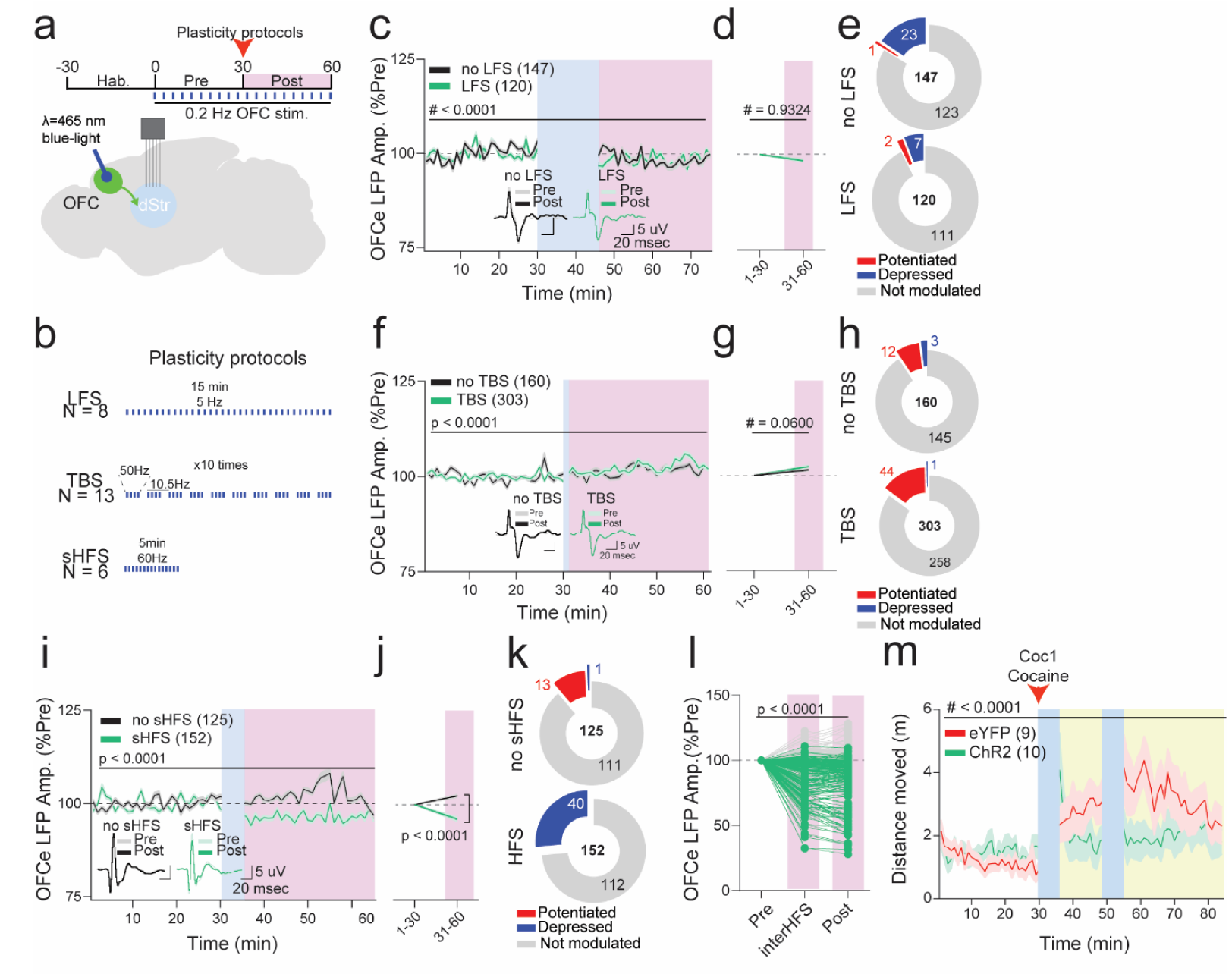
Characterization of HFS and other OFC-DMS stimulation protocols. (a) Brain schematic and experimental paradigm. (b) Schematics of plasticity protocols. (c) Time-course of normalized OFCe LFP response amplitude upon LFS or no LFS (RM two-way ANOVA; time main effect: F_(18.83,4970)_ = 13.24, p < 0.0001; protocol main effect: F_(1,264)_ = 0.007208, p = 0.9324; time Χ protocol interaction: F_(59,15576)_ = 6.597, p < 0.0001). (d) Binned OFCe LFP amplitude for no LFS and LFS (RM two-way ANOVA: time main effect: F_(1,264)_ = 47.66, p < 0.0001; protocol main effect: F_(1,264)_ = 0.0072, p = 0.9324; time Χ protocol interaction: F_(1,264)_ = 0.0072, p = 0.9324; followed by Bonferroni post-hoc test no LFS vs LFS; 1-30: t_(528)_ = 0; 31-60: t_(528)_ = 0.1201). (e) Pie-charts reporting the number of non-modulated, potentiated and depressed OFCe LFP responses upon LFS or no LFS. (f) Time-course of normalized OFCe LFP response amplitude upon TBS or no TBS (RM two-way ANOVA; time main effect: F_(59,27660)_ = 13.50, p < 0.0001, protocol main effect: F_(1,27660)_ = 20.58, p < 0.0001; time Χ protocol interaction: F_(59,27660)_ = 5.518, p < 0.0001). (g) Binned OFCe LFP amplitude for no TBS or TBS (RM two-way ANOVA; time main effect: F_(1,922)_ = 65.84, p < 0.0001; protocol main effect: F_(1,922)_ = 3.545, p = 0.0600; time Χ protocol interaction: F_(1,922)_ = 3.545, p = 0.0600 followed by Bonferroni post-hoc test no TBS vs TBS; 1-30: t_(922)_ = 0, p > 0.9999; 31-60: t_(922)_ = 2.663). (h) Pie-charts reporting the number of non-modulated, potentiated and depressed OFCe LFP responses upon TBS or no TBS. (i) Time-course of normalized OFCe LFP response amplitude upon sHFS or no sHFS delivery (RM two-way ANOVA; F_(12.99,3571)_ = 9.344, p < 0.0001, protocol main effect: F_(1,275)_ = 81.42, p < 0.0001; time Χ protocol interaction: F_(59, 16225)_ = 21.99, p < 0.0001). (j) Binned OFCe LFP amplitude for no sHFS or HFS protocol (RM two-way ANOVA; time main effect: F_(1,275)_ = 5.083, p = 0.0250; protocol main effect: F_(1,275)_ = 81.42, p < 0.0001; time Χ protocol interaction: F_(1,275)_ = 81.42, p < 0.0001; followed by Bonferroni post-hoc test no sHFS vs sHFS; 1-30: t_(550)_ = 0, p > 0.9999; 31-60: t_(550)_ = 12.76). (k) Pie-charts reporting the number of non-modulated, potentiated and depressed OFCe LFP responses upon sHFS or no sHFS. (l) Binned OFCe LFP responses for HFS and no HFS (RM two-way ANOVA; time main effect: F_(2,792)_ = 91.45, p < 0.0001; protocol main effect: F_(1,396)_ = 207.2, p < 0.0001; time Χ protocol interaction: F_(2,792)_ = 184.3, p < 0.0001 followed by Bonferroni post-hoc test no HFS vs HFS; Pre: t_(1188)_ = 0, p > 0.9999; interHFS: t_(1188)_ = 16.64, p < 0.0001; Post: t_(1188)_ = 17.89, p < 0.0001). (m) Time-course distance moved at Coc1 (RM two-way ANOVA; time main effect: F_(73,1241)_ = 3.596, p < 0.0001; virus main effect: F_(1,17)_ = 2.210, p = 0.1555; time Χ virus interaction: F_(73,1241)_ = 1.728, p = 0.0002). Data are represented as single points and/or as mean ± SEM. N indicates number of units or animals included in the analysis.

## MATERIALS and METHODS

### Animals

Wildtype (WT), D1Cre (GENSAT: EY217) and A2aCre (GENSAT: KG139)^28^ mice on a C57Bl6/j background were used: 21 animals (12 males and 9 females) for *in vivo* electrophysiology experiments; 14 animals (9 males, 5 females) for fiber photometry, 20 animals (14 males and 6 females) for behavioral assessment of cocaine-induced hyperlocomotion and 8 males for phospho-c-Fos experiments. The animals were housed at the NIH research animal facility in standard vivarium cages with *ad libitum* food availability and 12-hour dark/light cycle. The experiments described here were conducted during light-period (typically between 9am and 7pm). All experimental procedures were approved by the National Institute of Diabetes and Digestive and Kidney Diseases/National Institutes of Health Animal Care and Use Committee.

### Viral infusions and optic fiber implantation

Viral infections of OFC and DMS were conducted on adult male and female mice (older than 12 weeks). Anesthesia was induced with isoflurane at 2-3% and maintained during the entire surgery with isoflurane at 0.5-1.5%, delivered *via* a mouse mask mounted on a stereotaxic apparatus (Stoelting). Ear bars and mouth holder were used to keep the mouse head in place while the skin was shaved and disinfected with a povidone/iodine solution. The skull was exposed and a hole of approximately 0.5-1 mm diameter was performed with a microdrill. A 5 μL Hamilton syringe was connected to a 33gauge steel injector (Plastics1) via a hydraulic system. The injector was pre-loaded with AAVs and gently lowered into the brain at the following coordinates: OFC: AP + 2.5 mm, ML +1.1 mm, DV −2.5 mm; DMS: AP +0.5 mm, ML +1.5 mm, DV −2.8 mm (from bregma). A total volume of 500 nl of viral solution was delivered at each infection site with a syringe pump (Harvard apparatus) at a rate of 50 nL/min. The injector was left in place for 5 minutes after the infusion.

For *in vivo* electrophysiology experiments, the animals received a unilateral infection of the OFC. After removal of injector, suture (Coated VICRYL, polyglactin 910; Ethicon) followed by povidone/iodine were applied to close the wound. Animals were placed back in their home-cages on a pre-heated pad at 37°C until complete recovery and subcutaneously injected with buprenorphine (0.05 mg/kg). Animals were allowed at least 2 weeks for viral expression before electrodes were implanted in a second surgery.

For fiber photometry experiments, immediately after unilateral viral infusion in DMS, a 5 mm long fiber optic cannulae (200 μm, 0.39 NA, 1.25 mm) with stainless steel ferrule (Thorlabs) was gently lowered into DMS between 0.5-0.3 mm above the infusion site for GCamP6s-emitted fluorescence recordings.

For behavioral experiments with cocaine and OFC-DMS input stimulation, mice received bilateral opsin viral infusions in the OFC. In the same surgery, two fiber optic cannulae (200 μm, 0.39 NA, 1.25 mm, ceramic ferrule) were lowered bilaterally into DMS for OFC terminal stimulation (AP +0.5 mm, ML ±1.5 mm, DV −2.8 mm).

Optic fibers were fixed to the skull with a layer of C&B Metabond® Quick Adhesive Cement System and the implant was fortified with acrylic dental cement. Upon solidification of the implant, animals were placed back in their home-cages on a pre-heated pad at 37°C. After full recovery, animals received a subcutaneous injection of buprenorphine (0.05 mg/kg).

### Array and optic fiber implant for *in vivo* electrophysiology recordings

Implants of recording arrays for *in vivo* extracellular recordings and OFC stimulation typically occurred two weeks after viral infusions. In this study, we used 32-channel offset micro-arrays composed of 35 μm tungsten electrodes with polished tips and spaced by 150 μm. Uncleaved fiber optic cannulae (200 μm, 0.50 NA, 1.25 mm) with ceramic ferrule were purchased from Thorlabs and cut at the desired length for OFC stimulation. After anesthesia and skull exposure, two holes were drilled at the stereotaxic coordinates reported above. The optic fiber tip was lowered 0.3 mm above the infusion site with a 10° angle in the OFC, while the array was lowered into DMS. A grounding wire was inserted into the parietal lobe and a mini-screw was placed contro-laterally to the array to stabilize the implant. Both the optic fiber and the array were fixed to the skull and screw with C&B Metabond® Quick Adhesive Cement System and then cement with dental acrylic. After surgery, animals were transferred to pre-heated pad until complete recovery and subcutaneously injected with buprenorphine (0.05 mg/kg). Mice were single-housed.

### Fiber photometry and calcium population activity analysis

About 2-4 weeks after surgery, mice (8 D1Cre and 6 A2aCre) were acclimated for 30 min/day at least twice to a Phenotyper box (Noldus, PT T10/N, 24 VDC – 0.6A) with dimensions 30 cm x 30 cm x 34 cm. On the experimental day, mice were recorded for 30 minutes, before receiving a saline or cocaine i.p. injection (20 mg/Kg; counterbalanced). A mating sleeve (zirconia) connected the stainless-steel ferrule to the patch-cord (200μm core optic fiber, 0.48 NA, Doric), which both transmitted excitatory blue-light (wavelength: 475 λm, power: ~30 μW) and collected GCaMP6s-emitted photons. An optical commutator (Doric) was used to allow for rotation of the mice without tangling. The emitted light passed through a dichroic mirror and 505-535 nm filter (FMC4 port minicube, Doric) and then measured with a photodetector (Model 2151, Newport). GCaMP6s signal was collected, digitized and measured with Omniplex acquisition system (Plexon, Inc). The change in fluorescence (dF) was normalized to total fluorescence (F) using a custom Python scripts run in NeuroExplorer (script available on request). This script calculated a moving window of 2 minutes around each data point and used this as F, sliding this window along the entire recording trace to normalize each recorded data point. This approach normalizes the data and corrects for bleaching in one step. Animal movements in the open field were tracked and analyzed with Noldus Ethovision software. Fluorescence events (>5% dF/F, with 1 sec-long inter-event interval) were identified and binned in 60 sec time intervals. A time-course was generated by normalizing 1-min bin frequency values to the average of 30 bins in the pre-injection period and expressed as percentage (% of Pre). Larger binned frequency values for pre- and post-injection periods were obtained by averaging the 1-min long bins within 11-30 min for Pre- and 41-60 min for post-injection periods.

### *In vivo* electrophysiology recordings

After recovery from surgery (between 7-10 days), mice were connected to the recording setup and acclimated to an open field, 34.5 cm (l) x 34.5 cm (w) x 34.5 cm (h), head-stage and cable for at least 4 sessions of 30 minutes each on different days. On the experimental day, multi and single-units were collected via an Omniplex neurophysiology system (Plexon Inc) through a multiplexing head-stage (Triangle Biosystems). Spike channels were acquired at 40 kHz with 16-bit resolution, and band-pass filtered at 150 Hz to 3 kHz before spike sorting, while local field potentials (LFP) were digitized at 1 kHz. Single and multi-units were sorted by principal component analysis performed with Offline sorter software and MANCOVA test determined significant clustering of single-units *vs* multi-units. To determine the identity of striatal neurons, we calculated the waveform duration (time peak-to-valley: MSN > 350 μsec; interneuron < 350 μsec).

### Pharmacological and optogenetic manipulations for *in vivo* electrophysiology recordings

Animals to pharmacological agents, optogenetic stimulation or both. Mice received intraperitoneal injections of psychostimulants (cocaine and amphetamine, and their respective controls), dopaminergic antagonists (SCH23390, Sulpiride and their respective vehicle controls) and GBR13069 (and its control). *In vivo* plasticity was tested *via* the delivery of blue-light pulses into the OFC at different frequencies: low-frequency stimulation (LFS) consisted in 15 minute-long delivery of 15msec-long blue-light pulses at 5 Hz; theta-burst stimulation (TBS) consisted in the repeated delivery (10 times) of 10 trains of action potentials (at theta-frequency: 10.5 Hz) organized in 4 light-pulses at 50 Hz; short-high frequency stimulation (sHFS) consisted in the delivery of 5 msec-long blue light pulses at 60Hz; while HFS consisted in the delivery of 2-blocks of sHFS, interleaved by a 14 minutes interval during which animals received 15 msec-long blue-light pulses at 0.2 Hz.

### Analysis of *in vivo* electrophysiology recordings

The firing frequency of multi- and single-units during pre- and post-manipulation period was calculated in 60-sec bins with Neuroexplorer software. Average firing frequencies for pre- and post-manipulation periods were calculated as average of 30 bins in pre- and 30 bins in post-manipulation periods. A *paired t-test* followed by Bonferroni correction (α = 0.05/n of comparisons) was performed between pre- and post-manipulation period (10-30 minutes bin) to determine whether the manipulation induced significant changes in firing frequencies. To determine the direction of the change the average firing frequency during the post-manipulation was expressed as percentage of the pre-manipulation, such that changes >100% identified an increase while changes <100% indicated a decrease in firing frequency.

For OFC-stimulation evoked (OFCe) firing frequency, animals were subjected to 15 msec long blue-light pulses at 0.2 Hz, 30 minutes before and 30 minutes after pharmacological or optogenetic stimulation. The power of the laser was adjusted between 8-30 mW to obtain reliable responses. To determine significant changes in neuronal activity, firing frequency was analyzed 25 msec before (OFF period) and 25msec after the beginning of OFC illumination (ON period). A *paired t-test* between trial-by-trial values of firing frequency was used to determine significant modulation by OFC stimulation. A *paired t-test* followed by Bonferroni correction between trial-by-trial (25 msec after the beginning of blue-light stimulation) pre-(1-30 mins) and post-manipulation (31-60 mins) periods was used to determine significant modulation in OFCe firing rates. A time-course of OFCe firing frequency was generated by binning OFCe firing (60 sec) and expressing it as percentage of the averaged OFCe firing during pre-manipulation period. Binned OFCe firing rate responses were obtained by averaging 60-sec bins for pre- and post-manipulation period.

For OFC-stimulation evoked local-field potentials (OFCe LFPs) responses, animals were subjected to 15 msec long blue-light delivery into OFC at 0.2Hz, 30 minutes before and 30 minutes after pharmacological or optogenetic manipulations. The power of the laser was adjusted between 8-30 mW to obtain reliable responses. OFCe LFP amplitude was measured by using a custom-made Python script as the difference between the minimum and a maximum deflection point within an interval of 100 msec after OFC stimulation. OFCe LFP amplitude was binned in 60-sec bins and expressed as percentage of averaged OFCe LFP amplitude during pre-manipulation period. Significantly modulated OFCe LFP responses were identified by paired t-test followed by Bonferroni correction (α = 0.05/n of comparisons). To determine the direction of the change of OFCe LFP responses, the averaged OFCe LFP amplitude during the post-manipulation period was expressed as percentage of the averaged OFCe LFP amplitude in pre-manipulation period, such that changes >100% identified an increase while changes <=100% indicated a decrease in firing frequency. Binned OFCe LFP responses were obtained by averaging 60-sec bins for pre- and post-manipulation period.

Animal movement in the arena was tracked via Omniplex system software Cineplex. X and Y coordinates were used to determine instant velocity (1 sec). Instant velocity was averaged and binned (60 sec) and extracted via Neuroexplorer. Binned velocity in post-manipulation period was then normalized to the averaged binned velocity in pre-manipulation period, to generate time-course of normalized velocity (expressed as a percentage of Pre).

### Behavioral assay of sensitization to cocaine

The sensitization of psychomotor responses to cocaine was tested in an open-field box of 30 cm (l) x 30 cm (w) x 43 cm (h). eYFP or ChR2-expressing animals were first habituated to the novel environment and patch-cord for 3 consecutive days. On habituation days, animals were acclimated to the box for 30 minutes before receiving an i.p. injection of saline (without any optical stimulation). Starting from day 4, animals were acclimated to the box for 30 minutes before receiving an i.p. injection of cocaine (20 mg/Kg) immediately followed by HFS, with a laser power of 8mW measured at the tip of the optic fiber. This protocol was repeated for 4 more consecutive days, for a total of 5 days of cocaine+HFS. After 10 days of withdrawal, animals were acclimated to the box for 30 minutes and received an i.p. injection of cocaine without any optic stimulation. Animal’s activity (distance moved and locomotion) was tracked with Ethovision software. Distance moved was calculated as the sum of binned distance moved during the 14 minutes inter-HFS and 16 minutes post-HFS. We excluded from the analysis a ChR2-expressing animal that at day 1 showed a distance moved higher that 3 standard deviations from the group mean.

### Drugs

Cocaine hydrochloride (20mg/Kg; NDC 51552-0881-1, Fagron) and d-amphetamine hemisulfate salt C-IIN (3 mg/Kg; A5880, Sigma) were obtained via NIH pharmacy and dissolved in saline (NaCl 0.9%) to perform intra-peritoneal (i.p.) injection of 200-300 μL. (S)-(-)-Sulpiride (25 mg/Kg; 0895, Tocris), SCH23390 (0.15 mg/Kg; 0925, Tocris) and GBR13069 dihydrochloride (20 mg/Kg; 0420, Tocris) were diluted in a vehicle solution of 10% DMSO in saline for i.p. injection.

### Viruses

rAAV2/Syn-Chronos-GFP (2.1×10^12^ vg/mL), rAAVDJ/PAAV-Ef1a-DIO-GCaMP6s (3.9×10^12^ vg/mL) and rAAV2/hSyn-eYFP (3.4×10^12^ vg/mL) were purchased from Virus Vector Core at University of North Carolina at Chapel Hill. AAV5.CAG.hChR2(H134R)-mCherry.WPRE.SV40 (Addgene20938M, 3.9×10^12^ GC/mL) was purchased from Addgene.

### Phospho-fos experiments, immunostaining and cell counting

Mice (5 D1-tmt and 3 D2-gfp) were sacrificed 2 hours after an i.p. injection of cocaine (20 mg/kg) or saline (n=4 mice each). Brains were extracted and sectioned at 40 μm on a vibratome (Precisionary Compressotome). Tissue slices containing the striatum were immunostained for phospho-c-Fos (Cell Signaling monoclonal antibody #5348), with fluorescent secondary antibodies (Alexa 488 for D1-tmt mice and Alexa 555 for D2-gfp mice).

For quantification of striatal expression area, 2 striatal hemispheres per mouse were imaged on a scanning epifluorescence microscope (Leica DM6B). Phospho-c-Fos positive nuclei were identified manually in ImageJ and their X and Y coordinates were exported, registered to a common Atlas space, and plotted as heatmaps with MatPlotLib in Python 3.7. For colocalization with D1-tmt or D2-gfp, six two-color 10x fields of view were acquired from the DMS for each mouse on a confocal microscope (Leica microsystems). Images were quantified in ImageJ using the CellCounter plugin, where each phospho-c-Fos positive nucleus was also scored as positive or negative for the complementary fluorophore for cell type identification. To quantify the relative co-localization with D1R and D2R-expressing MSNs, confocal images were only acquired if they included multiple Fos positive nuclei (mean = 16.2, range = 3 to 32 Fos positive nuclei. This approach over-sampled Fos positive nuclei in the images from in the saline group. We therefore normalized the number of co-labeled neurons by the total Fos in the striatum of each mouse to obtain the extrapolated total counts in Figure 2e.

### Histological verification of sites of implant

At the end of the experiments, we performed a histological verification of implant placement. Animals were anesthetized (Chloral Hydrate, 7%) and perfused with 4% formaline. After over-night incubation in a 30% sucrose solution, either coronal or sagittal brain slices containing OFC and DMS were prepared. Slices were mounted on microscope slides with a mounting media with Fluoromount-G^TM^, with DAPI (Invitrogen) and imaged with a confocal microscope (Zeiss). For *in vivo* electrophysiology, behavioral and fiber photometry experiments implant placement was assessed *via* observation of implant tract or electric-lesions (performed with a 5 sec-long pulse of 10 mA; Ugo Basile Lesion Making Device).

### Statistical analysis

The number of animals included in this study was chosen based on those used in similar publications. Shapiro-Wilk test was used to assess normality of two-sample distributions. If violated, Mann-Whitney or Wilcoxon matched-pairs signed rank tests were applied, otherwise unpaired or paired t-tests were used. For comparing multiple sample distributions, normality of sample distribution was assumed and one-way ANOVA, repeated-measures (RM) ANOVA or RM two-way ANOVA were applied. When sphericity was not assumed, Geisser-Greenhouse correction was applied. If main effects and/or interaction were significant, Bonferroni post-hoc tests were used to determine significant differences across sample distributions. In all tests, statistical significance was determined when p < 0.05. Data are expressed as mean ± SEM. GraphPad Prism 8 software was used for graph presentation and statistical analysis. Statistical analysis is reported in figure legends. On graphs, p refers to p-values of main effects, comparison between two-sample distributions or post-hoc tests, while # refers to p-values of two factor interaction upon RM or regular two-way ANOVA. Animals were randomly assigned to each experimental condition but experimenters were not blinded to the treatment conditions as stimulants produced obvious behavioral changes upon injection.

### Data and code availability

The data that support the findings of this study are available from the corresponding author upon reasonable request.

## ACKNOWLEDGEMENTS

We thank Wambura Fobbs, Bridget Matikainen-Ankney, Meaghan Creed, Kelly Tan, Anatol Kreitzer, and Camilla Belone for helpful discussions about this manuscript, and Maya Bluitt for assistance with immunostaining and confocal imaging. We thank the HHMI GENIE project for GCaMP reagents.

## AUTHOR CONTRIBUTIONS

SB performed electrophysiology, fiber photometry experiments and data analysis; NM performed electrophysiology, fiber photometry and behavioral experiments. SB and AK designed the study and wrote the manuscript.

